# Targeting G_i/o_ protein-coupled receptor signaling blocks HER2-induced breast cancer development and enhances HER2-targeted therapy

**DOI:** 10.1101/2021.06.15.448534

**Authors:** Cancan Lyu, Yuanchao Ye, Maddison M Lensing, Kay-Uwe Wagner, Ronald J. Weigel, Songhai Chen

**Affiliations:** The Departments of Neuroscience and Pharmacology, Roy J. and Lucille A. Carver College of Medicine, University of Iowa, Iowa City, Iowa; The Department of Oncology, Wayne State University School of Medicine, Detroit, Michigan; The Department of Surgery, Roy J. and Lucille A. Carver College of Medicine, University of Iowa, Iowa City, Iowa

## Abstract

G protein coupled receptors (GPCRs) are among the most desirable drug targets for human disease. Although GPCR dysfunction drives the development and progression of many tumors including breast cancer (BC), targeting individual GPCRs has limited efficacy as a cancer therapy because numerous GPCRs are activated. In this study, we sought a new way of blocking GPCR activation in HER2^+^-BC by targeting a subgroup of GPCRs that couple to G_i/o_ proteins (G_i/o_-GPCRs). Using cell lines and transgenic mouse models, we showed in mammary epithelial cells, HER2 hyperactivation altered GPCR expression, particularly, G_i/o_-GPCRs. G_i/o_-GPCR stimulation transactivated EGFR and HER2, which in turn activated the PI3K/AKT and Src pathways. Uncoupling G_i/o_-GPCRs from cognate G_i/o_ proteins by pertussis toxin (PTx) inhibited BC cell proliferation and migration *in vitro* and suppressed HER2-driven tumor formation and metastasis *in vivo*. Moreover, targeting G_i/o_-GPCR signaling via PTx, PI3K, or Src inhibitors enhanced HER2-targeted therapy. These results indicate that HER2 hyperactivation in BC cells drives aberrant G_i/o_-GPCR signaling, and G_i/o_-GPCR signals converge on PI3K/AKT and Src signaling pathways to promote cancer progression and the development of resistance to HER2-targeted therapy. Our findings suggest a new way to pharmacologically deactivate GPCR signaling to block tumor growth and enhance therapeutic efficacy.

## Introduction

Breast cancer is the most common cancer and the second-most common cause of cancer death in US women. Approximately 15% to 20% of all breast cancers overexpress ErbB2/HER2, and so are classified as HER2^+^ subtypes, which are associated with aggressive cancers with poor clinical outcomes (1). HER2 is a member of the ErbB family, which includes EGFR/ErbB1, ErbB2/HER2, ErbB3, and ErbB4—all transmembrane receptor tyrosine kinases (RTKs) (2, 3). ErbB2/HER2 has no known ligands but can homodimerize or heterodimerize with EGFR or HER3 (4). Dimerized HER2 activates a complex cascade of downstream signaling that primarily consists of the phosphatidyl inositol 3 Kinase (PI3K)/AKT and the mitogen-activated protein kinase (MAPK) pathways (4). Hyperactivation of HER2 induces breast tumor formation, progression and metastasis.

The most successful treatment for HER2^+^ breast cancer is HER2-targeted therapy (5). Several Food and Drug Administration (FDA)-approved anti-HER2 drugs, including the humanized monoclonal antibody, trastuzumab, and the small-molecule dual inhibitor of HER2 and EGFR, lapatinib, significantly improved clinical outcomes of patients with HER2^+^ breast cancer. Nevertheless, tumors that initially respond to HER2-targeted therapy can eventually develop resistance (5). Thus, to improve clinical outcomes of advanced HER2^+^ breast cancer, it is critical to develop novel therapeutic approaches that can improve efficacy of HER2-targeted therapy.

G protein coupled receptors are the largest family of cell surface receptors and consist of over 800 members that regulate a plethora of biological functions (6). GPCR dysfunction drives the development and progression of many tumors including breast cancer (7). Transcriptomic profiling shows breast cancer cells aberrantly express multiple GPCRs (8). In multiple molecular subtypes of breast cancer, proteogenomic analysis identifies aberrant GPCR activation (9).

In preclinical mouse models, diverse GPCRs (e.g., lysophosphatidic acid (LPA), thrombin, endothelin, prostaglandin E2 and many chemokine receptors, such as CXCR4 and CCR5, contribute to breast tumor growth and/or metastasis (10). Nevertheless, attempts to target individual GPCRs as a cancer therapy have failed in many clinical trials, due largely to the lack of efficacy (11). Indeed, although GPCRs are the most desirable drug targets and nearly 40% of currently marketed drugs target GPCRs, only a few are used in cancer (11). This failure is likely because many different GPCRs are dysregulated in cancer and have the same oncogenic effect. Thus, to harness the power of GPCRs for cancer treatment, alternative approaches must be developed to overcome the redundant nature of GPCRs in cancer. One approach to overcome this redundancy is to target the shared functions of the group of GPCRs that drive breast cancer.

Most GPCRs mediate cellular responses by activating heterotrimeric G proteins, which consist of Gα and Gβγ subunits (12, 13). Sequence homology among Gα subunits distinguishes four classes of G proteins: G_i/o_, G_s_, G_q/11_, and G_12/13_; and although GPCRs can activate more than one class, they prefer one class over another (12). Of the 376 non-sensory human GPCRs, approximately 111 are coupled to G_i/o_, 55 coupled to G_s_, 81 coupled to G_q/11_, 12 coupled to G_12/13_, and 153 have unknown G-protein linkage.

The G_i/o_-GPCRs appear to be particularly important in breast cancer. PTx, which catalyzes the ADP-ribosylation of the Gα_i/o_ subunits and selectively uncouples G_i/o_ proteins from their receptors (e.g., PAR1, LPA and chemokine receptors) (14), blocks the effects of most GPCRs implicated in cancer, particularly progression and invasion (15-18). Moreover, 2% of all breast cancers carry a constitutively active form of G_αo_ (R243H) that promotes oncogenic transformation of normal mammary epithelial cells (19, 20). Additionally, we and others previously showed that many G_i/o_-coupled GPCRs signals mediating breast tumor cell growth and migration *in vitro* and tumor growth and metastasis *in vivo* all converge at Gβγ (17, 21, 22). Thus, targeting G_i/o_-GPCRs may be an effective strategy for halting breast tumor progression and overcoming drug resistance.

In this study, we sought to determine the function of a whole set of G_i/o_-GPCRs in HER2-induced breast cancer development and assess whether targeting G_i/o_-GPCR signaling could augment HER2-targeted therapy. We show that aberrant Gi/o-GPCR signaling promotes breast cancer cell proliferation and migration *in vitro* and contributes to HER2-induced tumor development and metastasis in a genetically modified mouse model. Blocking G_i/o_-GPCR signaling also enhances the efficacy of the HER2-targeted therapy *in vitro* and *in vivo*. These findings demonstrate that targeting G_i/o_-GPCR signaling may represent a new approach to blocking tumor progression and augmenting HER2-targeted therapy.

## Results

### HER2 regulates GPCR expression in mammary epithelia

To identify the GPCRs that drive breast cancer progression, invasive breast cancers from the TCGA database were profiled for RNA expression of 376 non-sensory GPCRs. Each sample overexpressed several GPCRs that couple to G_i/o_, G_s_, G_q/11_, G_12/13_ or unknown G proteins (supplemental Figure 1A-E), but no single GPCR was overexpressed in all samples. However, since tumors contain mixed populations of cells, these analyses could not reveal the identity of cells that overexpress GPCRs.

Therefore, GPCR expression profiling was further done in tumors of a well-established transgenic animal model of human HER2^+^ breast cancer, Neu mice, which express an activated rat ErbB2/HER2 homologue selectively in the mammary gland (23). Mammary epithelia from the mammary gland of the wild-type *vs*. premalignant mammary tissue of age-matched, Neu mice were profiled for expression of 370 non-sensory GPCRs. The results showed 291 GPCRs were expressed by control and Neu epithelia, 264 expressed by both, 20 expressed in controls only and 7 GPCRs uniquely expressed in Neu cells (Figure. 1A). Of the 264 GPCRs expressed in both normal and cancer cells, most (106) have unknown G protein linkages (Figure. 1B). More of the remaining couple to G_i/o_ (80) than to other G proteins (Figure. 1B). As compared to the control, 133 GPCRs were upregulated more than two-fold in Neu epithelial cells (Table 1). Among these, 44 have unknown G protein linkages, 40 couple to G_i/o_, 28 couple to G_q_, 17 couple to G_s_, and 4 couple to G_12/13_ (Figure 1C, supplemental Figure 2 and Table 1). Of the G_i/o_-GPCRs differentially expressed in Neu mice, more were upregulated (22) than downregulated (18; Figure. 1C and table 1).

**Figure 1.**
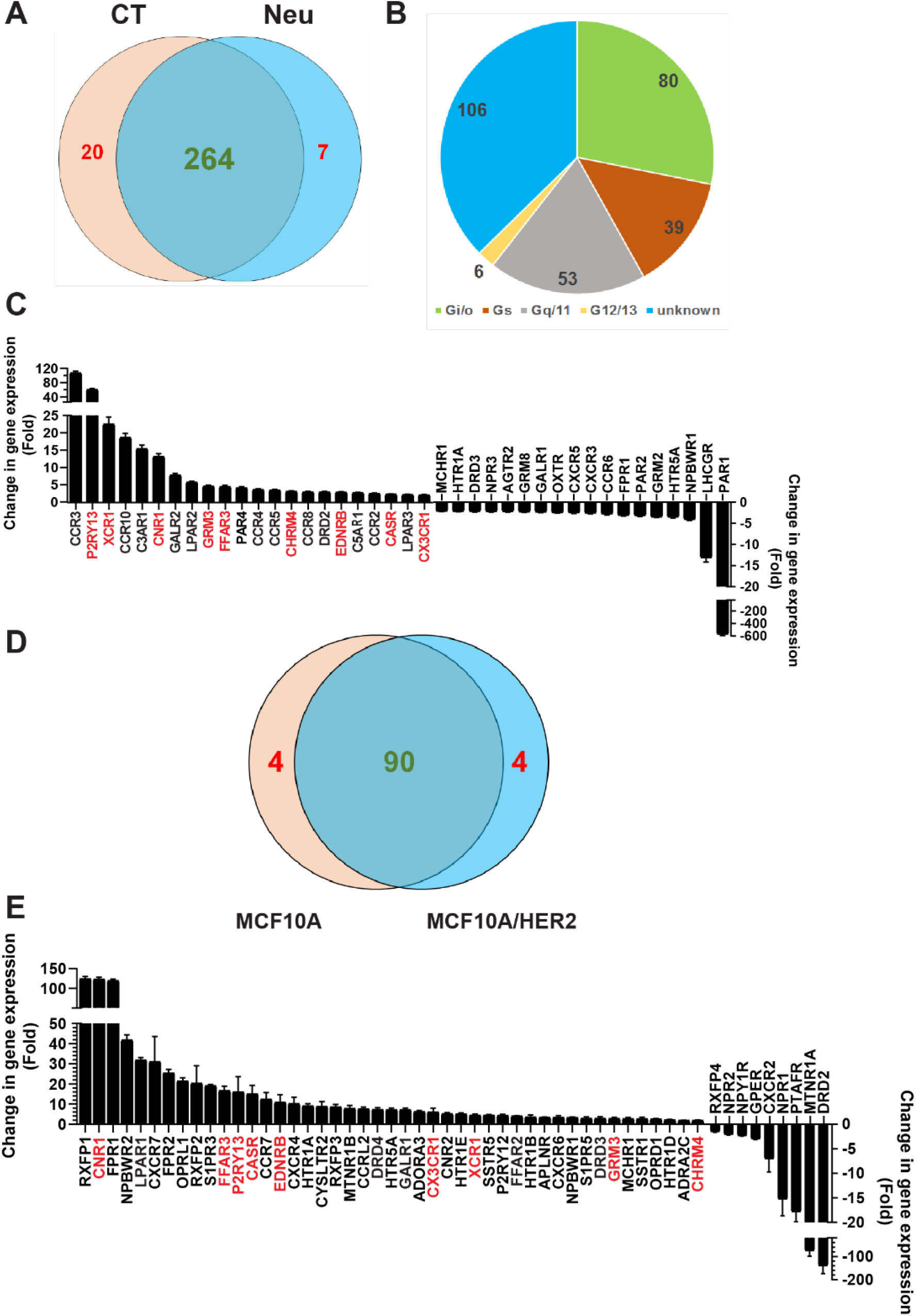
HER2 regulates GPCR expression in mammary epithelial cells. A, Venn diagram showing the number of GPCRs expressed in control and Neu cells. B, G-protein linkage of the GPCRs expressed in common, in control and Neu cells. C, The G_i/o_-GPCRs showing more than a 2-fold change of expression in Neu cells as compared to control cells, N=3. D, Venn diagram showing the number of G_i/o_-GPCRs expressed in MCF10A and MCF10A/HER2 cells. E, The G_i/o_-GPCRs showing more than a 2-fold change of expression in MCF10A/HER2 cells as compared to control cells, N=3.

**Table 1:** The number of GPCRs exhibiting a greater than two-fold change of expression in Neu epithelial cells as compared to the control

| GPCRs | Upregulated | Downregulated | Total |
| --- | --- | --- | --- |
| Gi/o-coupled | 22 | 18 | 40 |
| Gs-coupled | 7 | 10 | 17 |
| Gq-coupled | 13 | 15 | 28 |
| G12/13-coupled | 2 | 2 | 4 |
| Unknown | 23 | 21 | 44 |
| Total | 67 | 66 | 133 |

To test further if the altered GPCR expression is induced by HER2 overexpression and hyperactivation, MCF10A and MCF10A overexpressing HER2 (MCF10A/HER2) cells were profiled for expression of 106 G_i/o_-GPCRs. Both lines expressed 90 receptors and each line expressed four unique genes (Figure 1D). Compared to MCF10A cells, MCF10A/HER2 cells upregulated more receptors (45 upregulated more than 2-fold) than were downregulated (9 downregulated; Figure. 1E). Among the 45 GPCRs upregulated in MCF10A/HER2 cells, 9 (*i*.*e*., CNR1, FFAR3, P2RY13, CASA, EDNRB, CX3CR1, XCR1, GRM3 and CHRM4) were also upregulated in Neu cells (Figure 1C). These findings indicate that HER2 overexpression and hyperactivation in breast epithelial cells alters GPCR expression, in particular, Gi/o-receptor upregulation.

To validate the function of the upregulated Gi/o-GPCRs in MCF10A/HER2 cells, we chose to stimulate MCF10A and MCF10A/HER2 cells with LPA and SDF1α, because their cognate receptors, LPAR1 and CXCR4 and CXCR7 were upregulated at ∼32, 10 and 31 fold, respectively, in MCF10A/HER2 cells *vs*. MCF10A cells (Figure. 1E). The phosphorylation of AKT^S473^ and Src^Y416^ stimulated by LPA and SDF1α increased significantly in MCF10A/HER2 cells *vs*. MCF10A cells, while increased ERK phosphorylation was only observed by LPA stimulation (supplemental Figure 3A-B). Moreover, PTx treatment suppressed the AKT, Src, and ERK phosphorylation that was stimulated by LPA- and SDF1α-but not EGF (supplemental Figure 3C-E), suggesting G_i/o_ proteins, in particular, drive LPA- and SDF1α-stimulated signaling.

### G_i/o_-coupled receptor signaling promotes HER2-induced tumor growth and metastasis

To test whether G_i/o_-GPCRs drive HER2^+^ breast cancer proliferation, we determined the effect of PTx treatment on MCF10A and MCF10A/HER2 cell growth in Matrigel. In Matrigel, MCF10A cells formed small, round, and well-organized acini, whereas MCF10A/HER2 cells formed large and disorganized colonies with multiple protrusions (Figure 2A-B). PTx treatment neither changed the size nor the number of MCF10A acini, but significantly reduced the size and number of MCF10A/HER2 colonies (Figure 2D-E). Similarly, treatment of Neu cells also decreased the size and number of mammospheres that grew in Matrigel (Figure 2C and 2F-G). The inhibitory effects of PTx on breast cancer cell proliferation could also be demonstrated in 2D-culture of several HER2^+^ breast cancer cell lines, including Neu, MCF10A/HER2, BT474, and BT474R, a trastuzumab-resistant BT474 derivative (Figure 2H-K). These findings suggest that G_i/o_-GPCR signaling contributes to HER2-induced mammary tumor cell growth but is dispensable for normal mammary epithelial cell growth.

**Figure 2.**
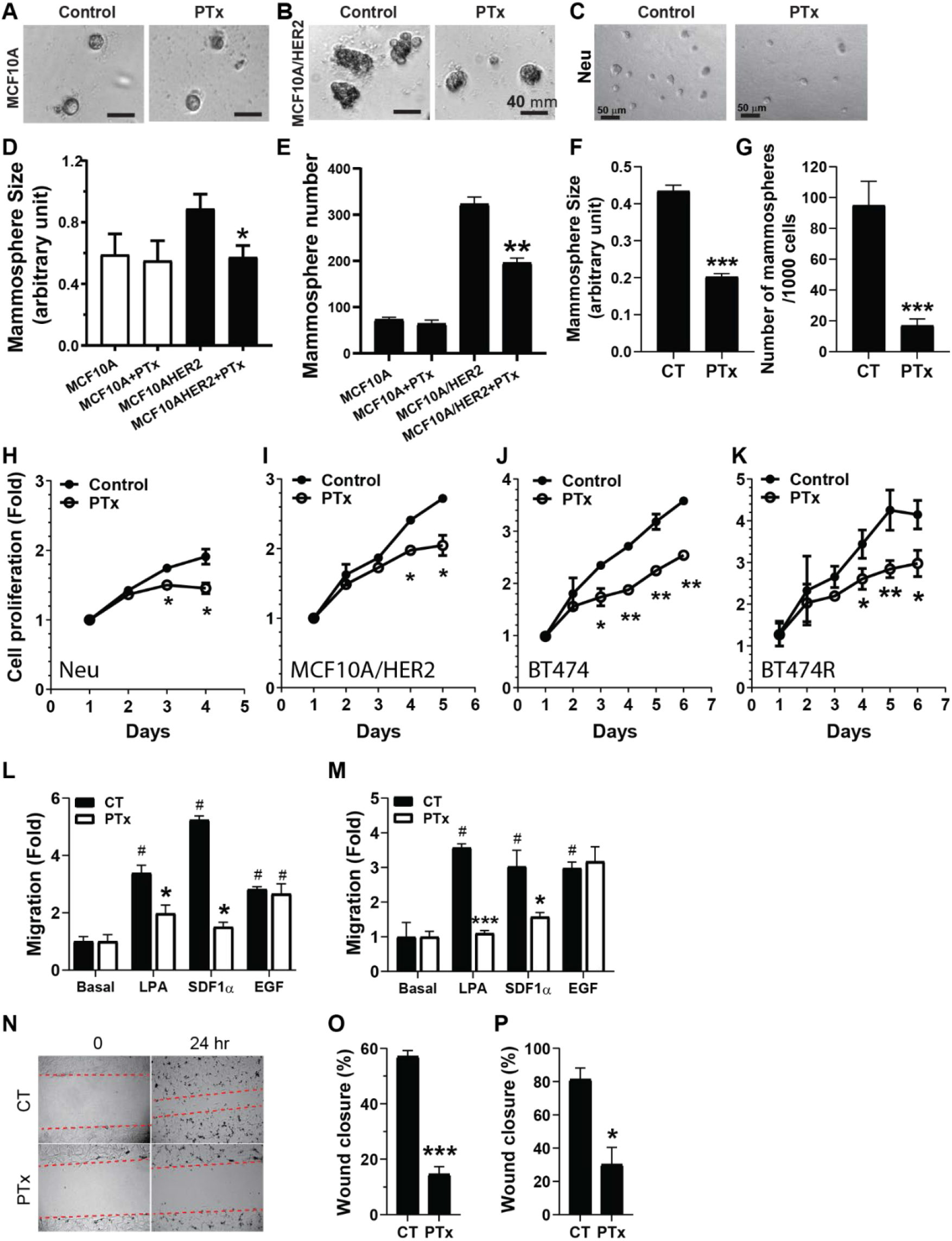
G_i/o_-GPCR signaling regulates the growth and migration of HER2^+^ mammary epithelial cells. A-G, representative images showing MCF10A (A), MCF10A/HER2 (B) and Neu (C) cells grown in Matrigel treated with vehicle control (CT) or PTx (0.2 μg/ml). D-G, quantitative data showing the size (D and F) and number of MCF10A (E and G), MCF10A/HER2 (D-E), and Neu (F-G) mammospheres. *p<0.05 *vs*. MCF10A/HER2 or CT, n=4. H-M, the effect of PTx treatment on the proliferation of Neu (H), MCF10A/HER2 (I), BT474 (J) and BT474R cells, and LPA-, SDF1α- and EGF-induced transwell migration of MCF10A/HER2 (L) and Neu (M) cells. *, **p<0.05 and 0.01 vs control (CT), n=4-6. N, representative images showing the size of the wound at 0 and 24 hours in MCF10A/HER2 cells treated with vehicle control (CT) or PTx. O-P, quantitative data showing the effect of PTx on wound healing in MCF10A/HER2 (O) and Neu (P) cells. *p<0.05 vs control (CT), n=6.

PTx treatment also decreased transwell migration of MCF10A/HER and Neu cells induced by LPA and SDF1α, but not by EGF (Figure 2L-M). In the wound-healing assay, MCF10A/HER and Neu cell migration was also inhibited (Figure 2N-P), indicating G_i/o_-GPCR signaling drives mammary tumor cell migration.

To corroborate these findings *in vivo*, we crossed the transgenic mice TetO-PTx (which carry a catalytic subunit of PTx under a tetO promoter) with the transgenic mice MMTV-tTA (which expresses the transactivator tTA from the mammary gland-specific MMTV promoter; Figure 3A). The resultant bi-genic offspring (tTA/PTx; TPTx) were crossed to Neu mice to generate tri-genics, tTA/Neu/PTx (Neu/PTx). qPCR analysis confirmed that PTx expression in their mammary glands was under doxycycline control (Figure 3B). Moreover, the PTx expression neither affected the litter size nor the survival of pups (data not shown); and whole-mount *in situ* staining of mammary glands did not detect a difference between mice with various genotypes in the number of terminal end buds (TNBs) at 1-month or the length of ductal distance in the mammary glands at different ages (supplemental Figure 4A-C), suggesting PTx did not affect mammary gland development. PTx expression, however, significantly delayed Neu-induced mammary tumor formation (161 days vs 190 days) and reduced tumor growth (Figure 3C-D). Ki67 staining showed that the percentage of Ki67^+^ tumor cells was significantly less in Neu/PTx tumors than Neu tumors (16.14 ± 2.99 vs 8.41 ± 2.0, p<0.05), suggesting tumor cell proliferation was inhibited by PTx (Figure 3E). To validate the inhibition of Neu tumor growth by PTx, we isolated Neu and Neu/PTx tumor cells and implanted an equal number of them into FVB/N mice fed normal or doxycycline-containing chow (to control PTx expression). In mice fed normal chow, Neu cells grew larger tumors than Neu/PTx cells (Figure 3F). This difference in tumor growth was abolished in mice fed doxycycline-containing chow (which suppressed PTx expression; Figure 3F).

**Figure 3.**
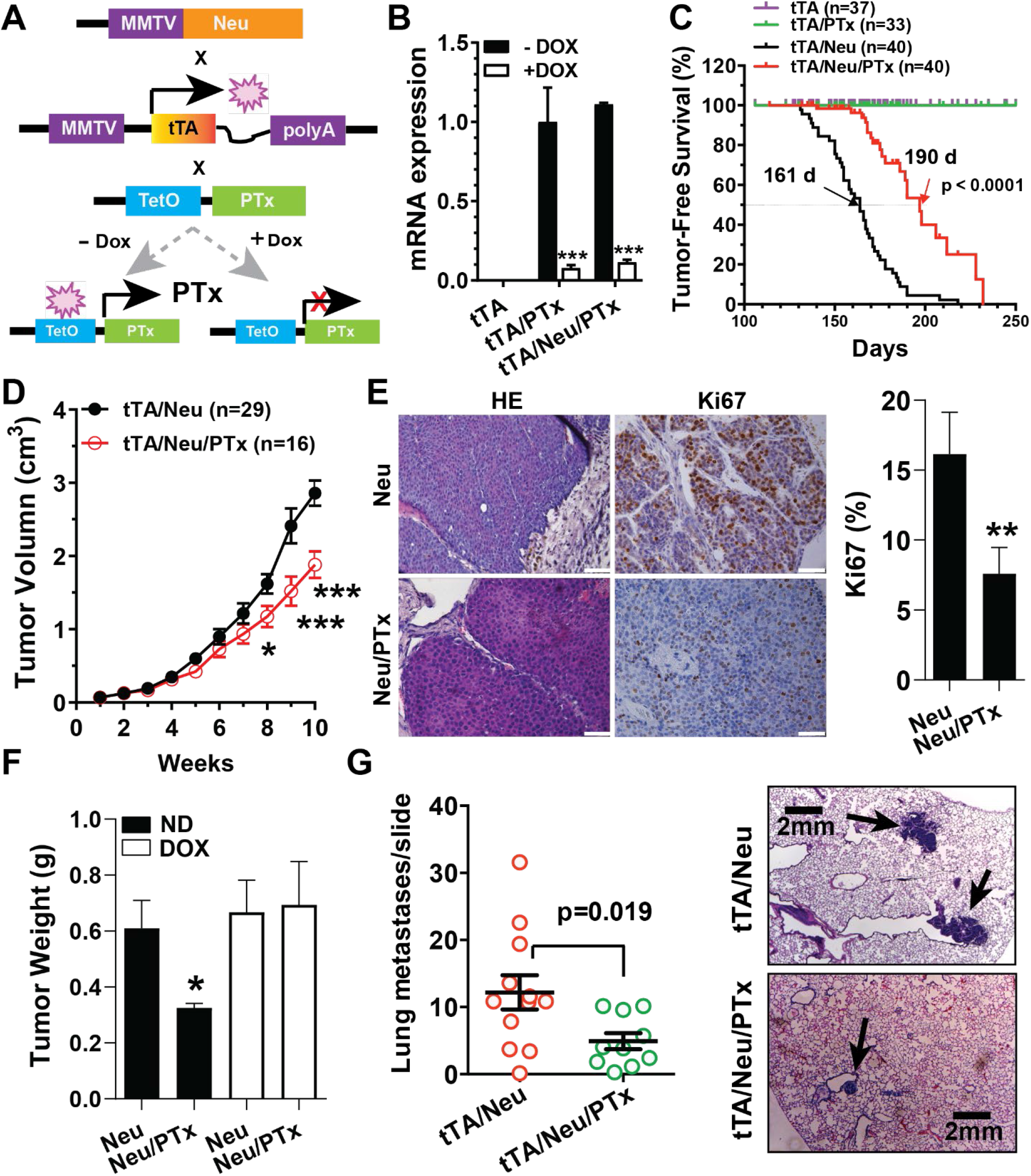
Gi/o-GPCR signaling contributes to HER2-induced mammary tumor development. A, schematic representation of mouse breeding and the regulation of PTx expression via doxycycline (Dox). B, qPCR results showing inducible PTx expression by doxycycline (DOX) in mammary epithelial cells. * p<0.05 *vs*. -Dox, n=4. C, tumor-free survival curves. D, Neu and Neu/PTx tumor growth curves. E, representative images showing HE and Ki67 staining of Neu and Neu/PTx tumors. Quantitatie data for Ki67 staining are shown in the right panel. Scale bar: 30 μm. **p<0.01 *vs*. Neu, n=9. F, the weight of tumors grown from Neu and Neu/PTx tumor cells in mice fed with normal chow (ND) or doxycyline-containing chow (DOX). *p<0.05 *vs*. Neu, n=7. G, the number of lung metastases and images representative of HE-stained lung metatases (indicated by arrows) from the transgenic Neu and Neu/PTx mice.

Lung metastases was checked when primary tumors reached a similar size (a diameter of ∼2.0 cm), showing that co-expression of PTx with Neu also reduced the size and number of lung metastases (Figure 3G). These findings indicate that G_i/o_-GPCR signaling contributes to HER2-induced tumor initiation, progression, and metastasis.

### G_i/o_-GPCRs crosstalk with EGFR and HER2

To identify mechanisms by which G_i/o_-GPCRs regulate HER2 tumor development and progression, we assessed the activation status of EGFR and HER2 and their downstream effectors. Immunoblotting found that, compared to Neu tumors, Neu/PTx tumors showed significantly reduced phosphorylation of EGFR and HER2, AKT^S473^ and Src^Y416^ (Figure 4A-B). The results were validated by IHC analysis (supplemental Figure 5A), suggesting that G_i/o_-GPCRs likely regulate EGFR and HER2 activation in tumor cells.

**Figure 4.**
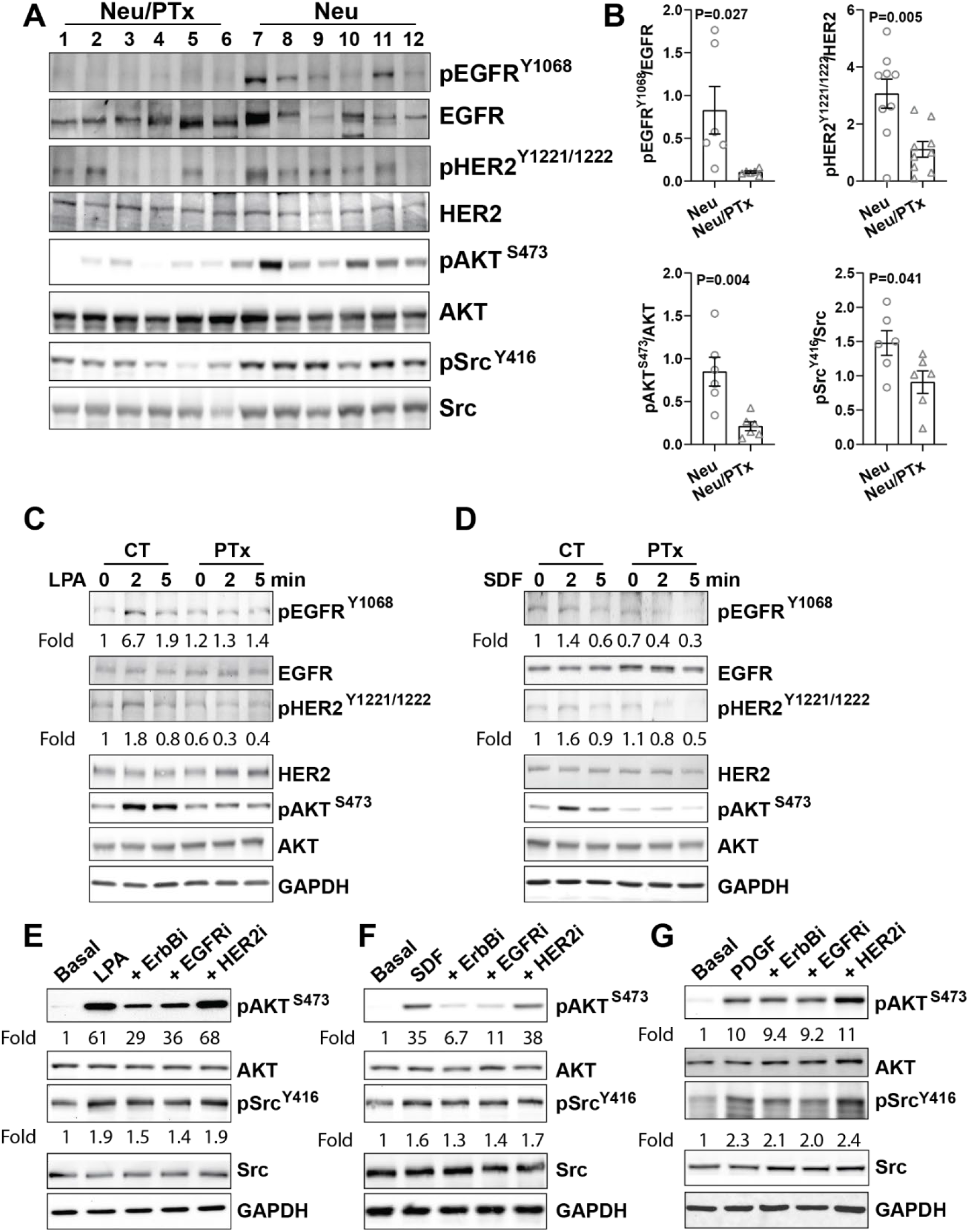
Gi/o-GPCRs induce transactivation of EGFR and HER2 in mammary tumor cells. A-B, Western blotting (A) showing decreased phosphorylation of EGFR^Y1806^, HER2^Y1221/1222^, AKT^s473^, and Src^Y416^ in Neu/PTx tumors as compared to Neu tumors. The Western blotting data were quantified and expressed as the ratio of the phosphorylated to total proteins (B). C-D, Western blotting showing phosphorylation of EGFR, HER2, and AKT in Neu cells treated with vehicle control or PTx and stimulated with LPA (C) or SDF1α (D). E-G, the effect of 1 μM of a pan-ErbB (sapitinib), EGFR (erlotinib)-, or HER2 (cp-724714)-specific inhibitor on LPA-(E), SDF1α-(F) or PDGF-(G) stimulated AKT and Src phosphorylation in Neu cells. The phosphorylation of EGFR^Y1068^, HER2^Y1221/1222^, AKT^S473^ and Src^Y416^ was quantified as the ratio of the phosphorylated to total proteins and expressed as the fold increase over the basal, which is indicated underneath the images.

To test if G_i/o_-GPCRs regulate EGFR and HER2 activities via transactivation, Neu cells were stimulated with LPA or SDF1α and then probed for EGFR and HER2 phosphorylation. As shown in Figure 4C-D, LPA and SDF1α stimulated EGFR and HER2 phosphorylation, which was abolished by PTx, indicating the involvement of G_i/o_ proteins. Similar results were found in MCF10A/HER2 cells (supplemental Figure 5B-C) and BT474 cells, which endogenously express HER2 (data not shown).

Next, we assessed whether EGFR and HER2 transactivation by G_i/o_-GPCRs drives activation of the shared effectors of G_i/o_-GPCRs and EGFR/HER2, such as AKT and Src. Neu and MCF10A/HER2 cells were treated with a pan-ErbB, EGFR-, or HER2-specific inhibitor, sapitinib (24), erlotinib (25), or cp-724714 (26); and the specificity of these inhibitors was confirmed in Neu cells stimulated with EGF. As expected, EGF stimulated phosphorylation of EGFR and HER2, was largely abolished by the pan-ErbB and EGFR-specific inhibitors, Sapitinib and erlotinib (supplemental Figure 5D). In contrast, the HER2-specific inhibitor, cp-724714 only inhibited EGF-stimulated HER2 phosphorylation, consistent with the idea that HER2 heterodimerizes with activated EGFR to form a signaling complex (supplemental Figure 5D). As shown in Figure 4E-F and supplemental Figure 5E-F, LPA- and SDF1α-stimulated AKT phosphorylation was partially suppressed by sapatinib and erlotinib, in both Neu and MCF10A/HER2 cells, while Src phosphorylation was only partially inhibited in Neu cells but not in MCF10A/HER2 cells. The HER2-specific inhibitor did not affect LPA- and SDF1α-stimulated AKT and Src phosphorylation, in either cell line. The effects of these inhibitors on LPA- and SDF1α-stimulated signaling are likely specific, since they had no effect on PDGF-stimulated AKT and Src phosphorylation in Neu cells (Figure 4G). These findings suggest that transactivation of EGFR by G_i/o_-GPCRs contributes to the activation of selective effectors, in a cell type-dependent manner.

### Targeting G_i/o_-GPCR signaling by PTx enhances the HER2-targeted therapy

Trastuzumab and lapatinib are the major anti-HER2 therapeutic reagents for HER2+ breast cancer (5). Since blocking G_i/o_-GPCR signaling suppressed HER2-induced tumor growth, we further tested if targeting G_i/o_-GPCRs affects the therapeutic efficacy of trastuzumab and lapatinib in Neu, MCF10A/HER2, BT474, and BT474R cells. After 5 days of treatment, trastuzumab significantly inhibited BT474 cell proliferation in a dose-dependent manner but had little effect on the growth of the other cells tested (Figure 5A, C and E). Both BT474 and Neu cells were sensitive but BT474R and MCF10A/HER2 cells were relatively resistant to lapatinib (Figure 5B, D, F and G). Notably, co-treatment with PTx enhanced the potency and/or efficacy of trastuzumab and lapatinib in all cells tested (Figure 5 and table 2).

**Figure 5.**
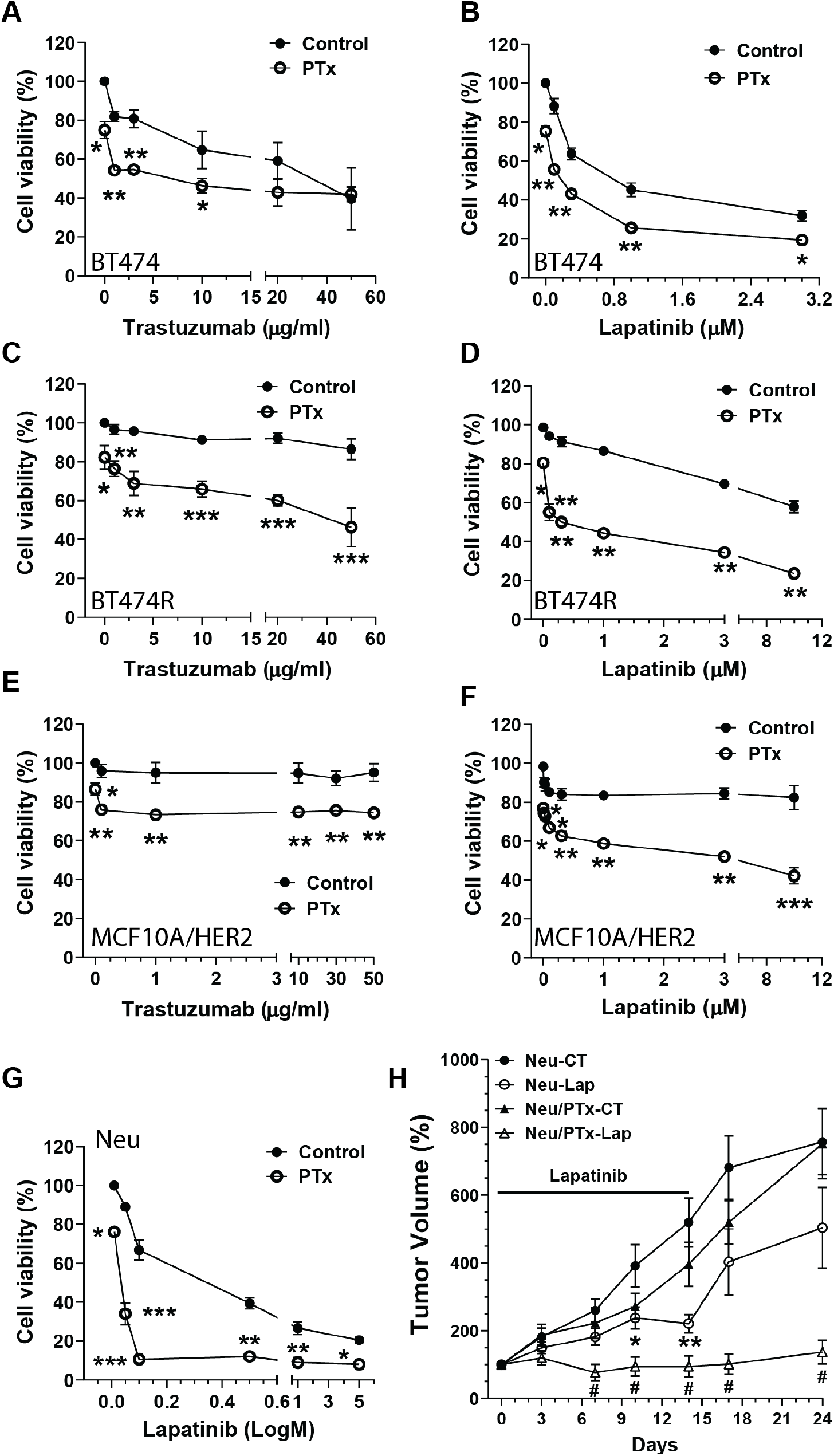
Targeting G_i/o_-GPCR signaling by PTx enhances HER2-targeted therapy. A-G, the effect of PTx (0.2 μg/ml) on the viability of BT474 (A-B), BT474R (C-D), MCF10A/HER2 (E-F) and Neu (G) cells treated with varying concentrations of trastuzumab for 5 days or lapatinib for 3 days. *, **, ***p<0.05, 0.01 and 0.001 vs control (CT), respectively, n=3-9. H, the growth curves of Neu and Neu/PTx tumors treated with vehicle control (CT) or lapatinib (Lap; 200 mg/kg, daily gavage). *, ** p<0.05 and 0.01 vs Neu-CT, respectively; # p<0.05 vs Neu-Lap. N=6-9.

**Table 2:** The IC_50_ and efficacy (E_max_) of the inhibitors in control and PTx-treated HER2^+^ mammary epithelical cells. E_max_ was determined as the maximum percentage inhibition at the highest concentrations of the inhibitors used. ND: undeterminable. Data are expressed as mean ± SEM. *, **, ***p<0.05, 0.01 and 0.001 vs control, respectively, n=3-9.

| Cells |  | Control |  | PTx |  |
| --- | --- | --- | --- | --- | --- |
|  |  | IC<sub>50</sub> | E<sub>max</sub> | IC<sub>50</sub> | E<sub>max</sub> |
| Trastuzumab | BT474 | 29.77 ± 1.46 | 61.5 ± 16.0 % | 4.85 ± 1.49*** | 59.1 ± 3.88 % |
|  | BT474R | ND | 13.5 ± 5.4 % | 89.57 ± 2.32 | 53.7 ± 9.9 %* |
|  | MCF10A/HER2 | ND | 4.9 ± 1.5 % | ND | 15.7 ± 2.1 %*** |
| Lapatinib | BT474 | 0.86 ± 0.11 | 68.1 ± 2.77 % | 0.088 ± 0.0122*** | 80.7 ± 1.89 %** |
|  | BT474R | 15.8 ± 1.2 | 42.2 ± 3.2 % | 0.26 ± 0.12*** | 76.5 ± 0.89 %*** |
|  | MCF10A/HER2 | ND | 17.6 ± 6.14 % | 10.79 ± 1.74 | 57.8 ± 4.18 %** |
|  | Neu | 0.261 ± 0.118 | 79.5 ± 0.9 % | 0.00847 ± 0.00121* | 92.0 ± 0.8 %*** |

To corroborate our findings *in vivo*, FVB mice were implanted with Neu and Neu/PTx tumors and maintained until the tumors grew to a comparable size (∼100 mm^3^); and for two weeks afterward, mice were treated by daily oral gavage with vehicle control or lapatinib (200mg/kg). Compared to the control, lapatinib partially suppressed the growth of Neu tumors, but completely blocked growth and even caused regression of Neu/PTx tumors (Figure 5H). Once the lapatinib treatment was stopped, Neu tumors resumed growth at an accelerated rate, while growth of the Neu/PTx tumors remained largely suppressed, over ten days (Figure 5H).

### Combination with PI3K and Src inhibitors enhances the HER2-targeted therapy

The PI3K/AKT and Src signaling pathways are critical for HER2-induced breast cancer progression, and resistance to HER2-targeted therapy (27-31). Since PI3K/AKT and Src are also the major pathways activated by many G_i/o_-GPCRs, we asked whether targeting PI3K/AKT and Src signaling mimicked the inhibitory effect of blocking G_i/o_-GPCRs by PTx. To do this, PI3K/AKT and Src signaling was blocked using a pan-PI3K inhibitor, GDC0941, and a Src-selective inhibitor, saracatinib. As shown in Figure 6A-C, these inhibitors selectively abolished EGF-, LPA-, and SDF1α-stimulated AKT, and Src phosphorylation in Neu cells, although saracatinib caused a slight inhibition of EGF-, LPA- and SDF1α-stimulated AKT phosphorylation, probably due to cross-talk between the Src and PI3K/AKT pathways.

**Figure 6.**
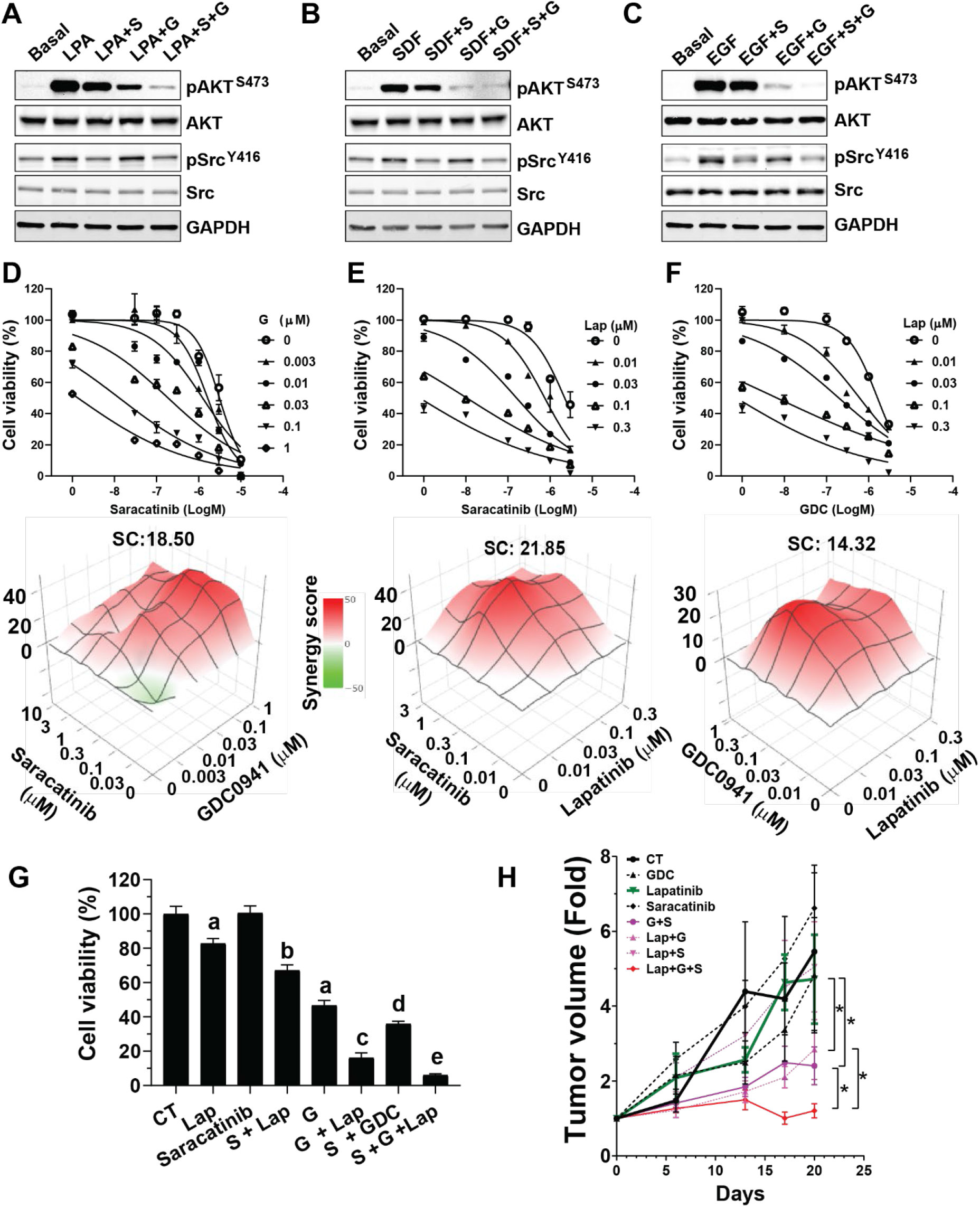
The combined effects of lapatinib plus PI3K and Src inhibitors on HER2_+_ breast cancer cells and tumors. A-C, effects of the PI3K and Src inhibitors, saracatinib (S) and GDC0941 (G), either alone or in combination, on LPA-(A), SDF1α-(B) and EGF-(C) stimulated AKT and Src signaling in Neu cells. D-F, effects of different drug combinations of saracatinib and GDC0941, saracatinib and lapatinib, and GDC and lapatinib on Neu cell growth. The dose-dependent inhibition curves and 3D plots for different drug combinations at varying concentrations are shown on the top and lower panels, respectively, The synergy score (SC) for each drug combination is also shown. G-H, effects of lapatinib (Lap), saracatinib (S), GDC0941 (G) and their combination on Neu cell (G) and tumor (H) growth. a, b, c, d, e indicate a statistically significant difference (p<0.05) from control (CT), Lap, Lap and G alone, S and G alone, S+G, respectively. *p<0.05, n=5-8.

When given alone, GDC0941 and saracatinib inhibited the proliferation of all breast cancer cells tested, in a dose-dependent manner (Figure 6D and Table 3). Checkerboard assays followed by the Bliss independent method analysis evaluated synergism between varying concentrations of GDC0942, saracatinib, and trastuzumab, or lapatinib. In Neu, MCF10A/HER, BT474, and BT474R cells, combining GDC0941 with saracatinib gave a significant synergistic effect (synergistic score >10; Figure 6D and Table 3). The combination of either GDC0941 or saracatinib with trastuzumab or lapatinib was also synergistic (Figure 6E-F and Table 3). Combining lapatinib with both saracatinib and GDC0941 inhibited Neu cell growth more than combining with either inhibitor alone (Figure 6G). Together, these findings suggest the PI3K and Src inhibitors enhance the therapeutic efficacy of trastuzumab and lapatinib.

The *in vivo* efficacy of combining the PI3K and Src inhibitors with lapatinib was tested against Neu syngeneic tumors. FVB mice were implanted with Neu cells, which were allowed to grow into tumors ∼100 mm^3^ in size. The mice were then treated by daily oral gavage with vehicle control, lapatinib (150mg/kg), GDC0941 (50 mg/kg) or saracatinib (10 mg/kg) alone or in combination, *i*.*e*., GDC0941 (50 mg/kg) + saracatinib (10 mg/kg), lapatinib (150mg/kg) + GDC0941 (50 mg/kg), lapatinib (150mg/kg) + saracatinib (10 mg/kg) or lapatinib (150mg/kg) + GDC0941 (50 mg/kg) + saracatinib (10 mg/kg), for three weeks. Under these conditions, lapatinib, GDC0941 or saracatinib alone, or lapatinib plus saracatinib did not significantly affect tumor growth (Figure 6H). Whereas a combination of GDC0941 with saracatinib or lapatinib partially suppressed tumor growth, strikingly, the combination of all three drugs not only blocked tumor growth but also caused tumor regression (Figure 6H).

## Discussion

Although multiple GPCRs are implicated in driving breast cancer formation and progression, the mechanisms underlying GPCR alteration in cancer are largely unknown (32). Moreover, to date, no approaches have effectively targeted multiple GPCRs as a cancer therapeutic (7, 11). Our studies show why this has been so difficult: in breast cancer cells, expression of a multiplicity of GPCRs is altered, a finding consistent with reports that various tumor types differentially express more than 50 GPCRs (8). Moreover, our data show HER2 overexpression can alter GPCR expression in both mouse and human mammary epithelia cells, suggesting HER2 signaling is a key regulator of GPCR mRNA expression. The upregulated GPCR expression likely contributes to aberrant GPCR signaling in tumors, since stimulation of the upregulated receptors, such as LPAR and CXCR4 and CXCR7 in MCF10A/HER cells, enhances AKT and Src signaling. Since constitutively active mutants of GPCRs are relatively rare in breast cancer (8), our results suggest GPCR upregulation by oncogenic signals may be the key mechanism contributing to aberrant GPCR signaling in tumors.

Our results are consistent with previous studies (8), and argue that Gi/o-coupled receptors are the main group of GPCRs that altered expression in breast cancer cells. Many members of the G_i/o_-GPCR family, such as LPAR, CXCR4, CXCR7, and PAR1, are known to play a role in breast cancer progression. However, targeting any one receptor as a cancer therapy has not achieved optimal efficacy in many clinical trials, likely because GPCRs function redundantly in promoting tumor progression. Our studies indicate that although not all G_i/o_-GPCRs are upregulated in tumor cells, the overall function of G_i/o_-GPCRs is to promote breast cancer formation and progression. This is demonstrated by findings that uncoupling G_i/o_-GPCRs to G_i/o_ proteins by PTx suppresses tumor cell growth and migration *in vitro*, and blocks tumor formation and progression *in vivo*. PTx effects are likely specific, as it selectively inhibits G_i/o_-GPCR-mediated but not EGF-stimulated signaling and does not affect normal mammary gland development. These findings further indicate that G_i/o_-GPCR signaling contributes to breast cancer growth and metastasis and may be targeted to block breast tumor progression. As a demonstration of the potential power of this approach for breast cancer therapeutics, we showed that blocking G_i/o_-GPCR signaling by PTx enhances the therapeutic efficacy of HER2-targeted therapy both *in vitro* and *in vivo*. Notably, blocking G_i/o_-GPCR signaling also sensitizes trastuzumab-resistant cells to trastuzumab, suggesting that targeting G_i/o_-GPCR signaling may represent a new approach to overcome resistance to HER2-targeted therapy.

PTx is a valuable tool for dissecting the function of G_i/o_-GPCRs, but it cannot be used as a therapeutic agent for blocking G_i/o_-GPCR signaling because it is a virulence factor of Bordetella pertusis and causes the respiratory disease pertussis (14). G_i/o_-GPCRs transmit signals through Gαi/o and Gβγ subunits; and signals originating from both are implicated in tumor progression, suggesting directly targeting G proteins may be an approach to blocking G_i/o_-GPCR signaling (6, 13). Yet, no Gαi/o-specific inhibitors are currently available (7). Although several small inhibitors of Gβγ have been developed, they may not selectively target G_i/o_-GPCR signaling since other classes of GPCRs can also initiate Gβγ signaling (33). Moreover, although Gβγ signaling is critical for growth and metastasis of triple-negative breast cancer cell lines (17, 21), whether it mediates progression of HER2-driven breast cancer remains unknown.

An alternative approach to targeting G_i/o_-GPCRs is to target their shared signaling pathways that are essential for cancer development and progression. Our results suggest PI3K/AKT and Src signaling represent such pathways, and so might be therapeutically targeted in HER2^+^ breast cancer. First, as reported here and previously (34-37), PI3K/AKT and Src can be activated downstream of multiple G_i/o_-GPCRs. The G_i/o_-GPCR-mediated mechanisms for PI3K/AKT and Src activation are complex and likely receptor-dependent. PI3K/AKT and Src may be activated by direct interaction with Gαi/o and Gβγ subunits or may be downstream of other signaling molecules such as small G proteins (34-38).

Second, PI3K/AKT and Src are key pathways activated by EGFR and HER2. PI3K/AKT and Src activities play key roles in HER2-induced breast tumor formation and progression (27-31). Moreover, they appear to be the convergence points for crosstalk between Gi/o-GPCRs and EGFR/HER2 (39-41). Although Gi/o-GPCRs induce transactivation of EGFR and HER2, the activation of EGFR and HER2 in turn contributes to Gi/o-GPCR-mediated PI3K/AKT and Src activation. Finally, several studies suggest that aberrant PI3K/AKT and Src activation contributes to resistance of HER2^+^ breast cancer to HER2-targeted therapy (31, 42-44). Consistent with these results, in breast cancer cell lines overexpressing HER2, the combination of PI3K or Src inhibitors with HER2-targeted therapy synergistically inhibits growth. Notably, recent reports demonstrate resistance to combined inhibition of PI3K and HER2 in HER2^+^ breast cancer is associated with upregulation of collagen, integrin β1 and Src activity in HER2+ breast cancer (45), suggesting that combined inhibition of both PI3K and Src might be superior to inhibition of the individual proteins in overcoming resistance to the HER2-targeted therapy. Our data support these findings and provide the first evidence that combined inhibition of both PI3K and Src synergize to block tumor growth and sensitize tumors to HER2-targeted therapy.

In conclusion, our study showed HER2 overexpression alters expression of multiple GPCRs in breasts cancer cells, and demonstrated that Gi/o-GPCRs, the largest subset of GPCRs, promote HER2-induced breast cancer formation and progression. Moreover, we provided a proof of concept that the Gi/o-GPCRs can be targeted, as a whole, to block tumor progression and enhance HER2-targeted therapy. Finally, our evidence shows PI3K and Src signaling represent the pathways shared downstream of Gi/o-GPCRs that may be targeted to enhance the efficacy of the HER2-targeted therapy.

## Methods

### Reagents

LPA and PTx were from Sigma. Human and mouse SDF1-α were from Pepro Tech. EGF from Gold Biotechnology. Sapitinib, erlotinib and cp-724714 were from Shelleck Chemicals. Trastuzumab was from Genetech. Laptinib, GDC0941 and saracatinib were from LC Laboratories. Antibodies for EGFR (no. 2232), phospho-EGFR^Y1068^ (no. 3777), HER2 (no. 2165), phospho-HER2^Y1221/1222^ (no. 2243), AKT (no. 4685), phospho-AKT^S473^ (no. 4060), Src (no. 2109), phospho-Src^Y416^ (no. 6943), ERK1/2 (no. 4696) and phospho-ERK1/2^T202/Y204^ (no. 4370) were from Cell Signaling Technology; GADPH (sc-47724) from Santa Cruz Biotechnology. Ki67 (GTX16667) from GeneTex. EGFR (no. 4267) from Cell Signaling Technology, and phospho-Src^Y418^ (ab4816) from Abcam were used in immunohistochemical staining.

### The Cancer Genome Atlas data analyses

The cBioportal for Cancer Genomics (www.cbioportal.org/) was used to analyze the mRNA expression of ∼400 non-sensory GPCR, in the breast invasive carcinoma dataset (TCGA, PanCancer Atlas). A two-fold z-score threshold identified patients with altered GPCR mRNA expression levels. The results are presented in heatmaps according to G protein linkage of GPCRs. The linkage of GPCRs to G proteins is assigned based on information for GPCRs in the IUPHAR/British Pharmacological Society (BPS) website.

### Cell lines

BT474 and MCF10A were purchased from the ATCC. The trastuzumab-resistant BT474 derivative (BT474R) and HER2-overexpressig MCF10A (MCF10A/HER2) cells were kindly provided by Dr. Hank Qi at the University of Iowa. Neu cells were generated from tumors arisen from the transgenic mice, MMTV-c-Neu, and cultured in DMEM media containing 10% FCS. Cell lines were tested for *Mycoplasma* using the Mycoplasma Detection kit (ATCC). BT474 and BT474R cells were cultured in DMEM/F12 media containing 10% FCS. MCF10A and MCF10A/HER2 were cultured in DMEM/F12 media containing 5% horse serum supplemented with EGF at 20ng/ml, hydrocortisone at 500ng/ml, cholera toxin at 100ng/ml and insulin at 10μg/ml. Each cell line was cryopreserved at low passage numbers (less than six passages after receipt) and used in experiments for a maximum of 18 passages.

### Isolation of mammary epithelial cells

Mammary epithelial cells were isolated from the mammary glands of 4-month-old wild-type and MMTV-c-Neu transgenic FVB/N mice, using the EasyStep Mouse Epithelial Cell Enrichment Kit (StemCell Technologies).

### Analysis of gene expression

Total RNA was extracted from the isolated mammary epithelial cells pooled from four mice, and MCF10A and MCF10A/HER2 cells using a Qiagen RNeasy MinitKit (Qiagen). cDNA was synthesized from 2 μg of total RNA using a High Capacity cDNA Reverse Transcription Kit (Applied Biosystems). The expression of 370, mouse, non-sensory GPCRs was analyzed using the RT2 Profiler PCR array (no. 330231) from Qiagen, as per the manufacturer’s instructions. 106 human Gi/o-GPCRs and the effects of PTx were analyzed with real-time qPCR using the SensiFAST SYBR No-ROX kit (Medidian Bioscience) and specific primers designed and synthesized by Integrated DNA Technologies (supplemental Table). Threshold cycle (C_t_) values of target genes were normalized to the mean C_t_ values of the housekeeping genes, actin, β2 microglobulin, Glyceraldehyde-3-phosphate dehydrogenase, β glucuronidase, heat shock protein 90 α. The expression of a gene was considered undetectable if its C_t_ value was <35. The fold changes of gene expression relative control are presented using the Graphpad Prism Software.

### Cell proliferation and viability assays

Cell proliferation in two-dimensional culture or in Matrigel was analyzed as we described previously (17, 22). To analyze the response of cells to drug treatment, 3000 cells were seeded in triplicate into a 96-well plate and varying concentrations of drugs were added the next day. Cells were exposed to trastuzumab for 5 days or other inhibitors for 3 days. Cell viability was quantified using AlamarBlue (Thermo Fisher Scientific) assays as per the manufacturer’s instructions. For combinational studies of two drugs, percentage inhibition was calculated and analyzed using the SynergyFinder 2.0 software (46).

### Cell migration and wound healing assays

Transwell migration and wound healing assays were performed as we described previously (17, 22). To exclude the influence of cell proliferation, cells were treated with 5 μg/ml mitomycin.

### Cell stimulation

Cells were seeded into 6-well plates. After 48 hours of serum starvation, cells were pre-treated with the indicated inhibitors for 1 hour, and then stimulated with various agonists for the indicated time. To determine the effect of PTx, cells were treated with PTx (200 ng/ml) for 24 hours.

### Western blotting analysis

Protein lysates were prepared from cells and tumor tissues and analyzed by Western blotting as we described, using the iBright CL1000 imaging system (Thermo Fisher Scientific) (17, 22).

### Mouse studies

MMTV-c-Neu mice were from Jackson Laboratory (no. 005038). TetO-PTx mice were from the Mutant Mouse Resource & Research Center (MMRRC; no. 014241) and MMTV-tTA mice were generated in Dr. Wagner’s laboratory (47). All mice were in the FVB/N genetic background. Mice were genotyped by PCR as reported previously (23, 47, 48). Female mice were kept as virgins throughout the experiments. To determine tumor onset, starting at four months after birth, mice were checked twice per week by palpation. To determine tumor progression, the largest tumor was measured weekly by caliper. To determine lung metastasis, the lung was harvested once the largest tumor reached a size of 2 cm in diameter, and was perfused and fixed with 4% paraformaldehyde before paraffin embedding. The number of metastases in the lung was analyzed by serial sectioning followed by HE staining.

To generate syngeneic tumor models, tumor cells were isolated from size-comparable tumors arisen from transgenic mice, i.e., MMTV-c-Neu/MMTV-tTA (Neu) and MMTV-c-Neu/MMTV-tTA/TetO-PTx (PTx), using the Mammary epithelial cell enrichment kit (StemCell Technologies). 1×10^6^ cells were injected into the right inguinal mammary fat pads of FVB/N mice. To determine the effect of PTx expression on tumor growth, mice were fed with normal chow or doxycycline-containing chow (625mg/kg; ENVIGO) to turn off PTx expression. Two months after turmor cells were implanted, tumors were excised and weighed. To determine the response of tumors to drug treatment, tumors were grown to a size of ∼100 mm^3^ and then treated with vehicle or drugs for 3 weeks. Tumor growth was monitored twice per week by caliper measurement.

### Whole-mount, histology and immunohistochemical analyses

Whole-mount staining of mammary glands from mice of different ages and histological and immunohistochemical analyses of tumors were performed as described (22, 49). Ki-67 was quantified by counting the number of positive nuclei for every 100 nuclei from multiple fields on the slide, to obtain the percentage of positive cells.

### Statistics

Data were expressed as mean ± SEM. Statistical comparisons between groups were analyzed by two tail Student’s *t* test or ANOVA (*P*<0.05 was considered significant).

### Study approval

All animal studies were conducted in accordance with an IACUC-approved protocol at the University of Iowa.

## Authors’ Contributions

Data curation, C. Lyu, and M. Lensing; Formal analysis, P C. Lyu, M. Lensing and S. Chen; Critical materials, K. Wagner; Methodology, P C. Lyu, and M. Lensing; Project administration, R. Weigel and S. Chen; Writing-original draft, S. Chen.; Writing-review & editing, R. Weigel and S. Chen.

## Achnowledgements

This work was supported in part by NIH grant R01CA207889 and DOD BCRP breakthrough award level 2 (BC151478), and an NCI Core Grant P30 CA086862 (University of Iowa Holden Comprehensive Cancer Center).

