## Supplemental Figures for "Targeting G_i/o_ protein-coupled receptor signaling blocks HER2-induced breast cancer development and enhances HER2-targeted therapy"

A

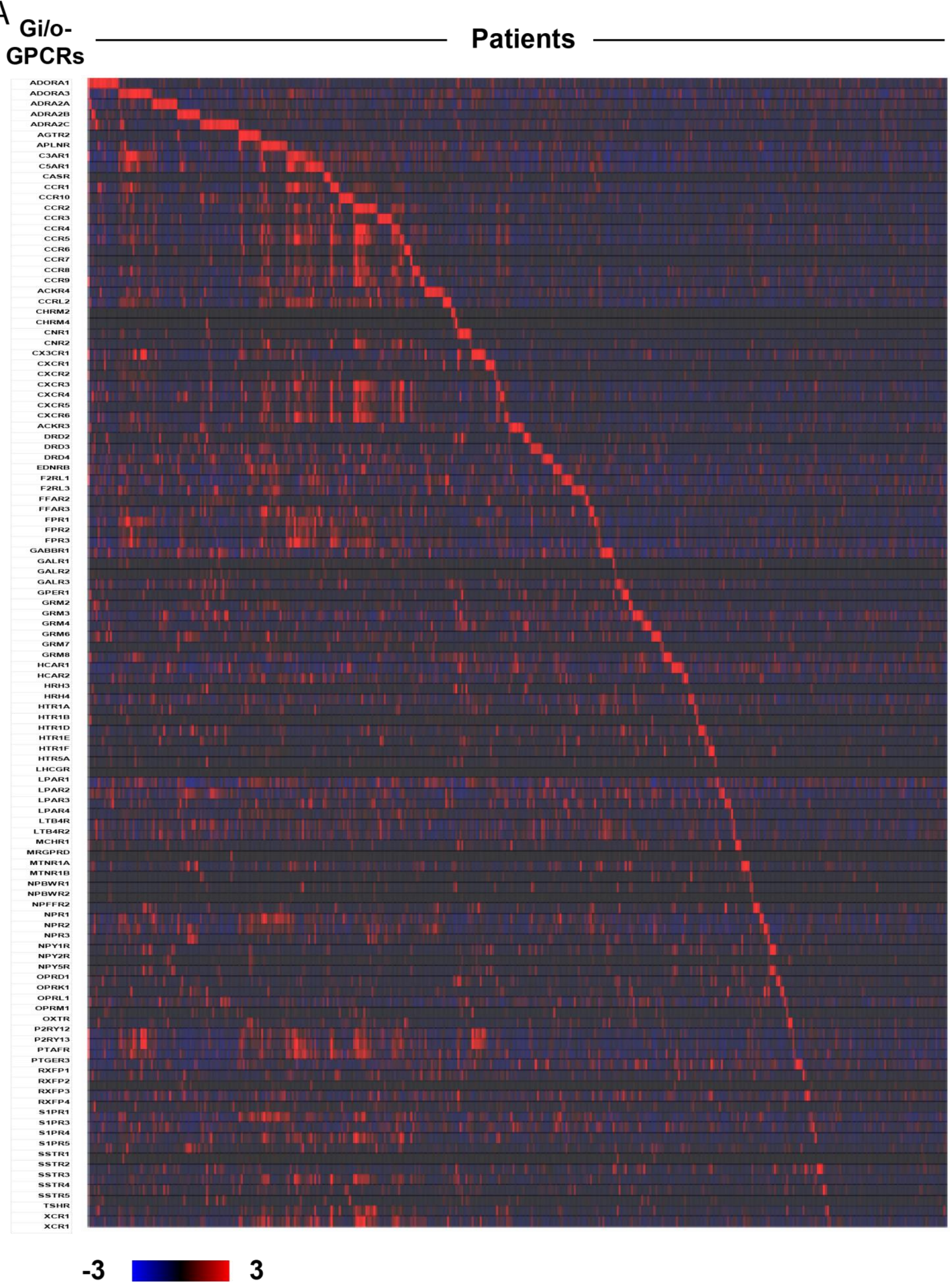
Gi/o-GPCRs
Patients
ADORA1
ADORA3
ADRA2A
ADRA2B
ADRA2C
AGTR2
APLNR
C3AR1
C5AR1
CASR
CCR1
CCR10
CCR2
CCR3
CCR4
CCR5
CCR6
CCR7
CCR8
CCR9
ACKR4
CCRL2
CHRM2
CHRM4
CNR1
CNR2
CX3CR1
CXCR1
CXCR2
CXCR3
CXCR4
CXCR5
CXCR6
ACKR3
DRD2
DRD3
DRD4
EDNRB
F2RL1
F2RL3
FFAR2
FFAR3
FPR1
FPR2
FPR3
GABBR1
GALR1
GALR2
GALR3
GPER1
GRM2
GRM3
GRM4
GRM6
GRM7
GRM8
HCAR1
HCAR2
HRH3
HRH4
HTR1A
HTR1B
HTR1D
HTR1E
HTR1F
HTR5A
LHCGR
LPAR1
LPAR2
LPAR3
LPAR4
LTB4R
LTB4R2
MCHR1
MRGPRD
MTNR1A
MTNR1B
NPBWR1
NPBWR2
NPFFR2
NPR1
NPR2
NPR3
NPY1R
NPY2R
NPY5R
OPRD1
OPRK1
OPRL1
OPRM1
OXTR
P2RY12
P2RY13
PTAFR
PTGER3
RXFP1
RXFP2
RXFP3
RXFP4
S1PR1
S1PR3
S1PR4
S1PR5
SSTR1
SSTR2
SSTR3
SSTR4
SSTR5
TSHR
XCR1
XCR1
-3
3

B

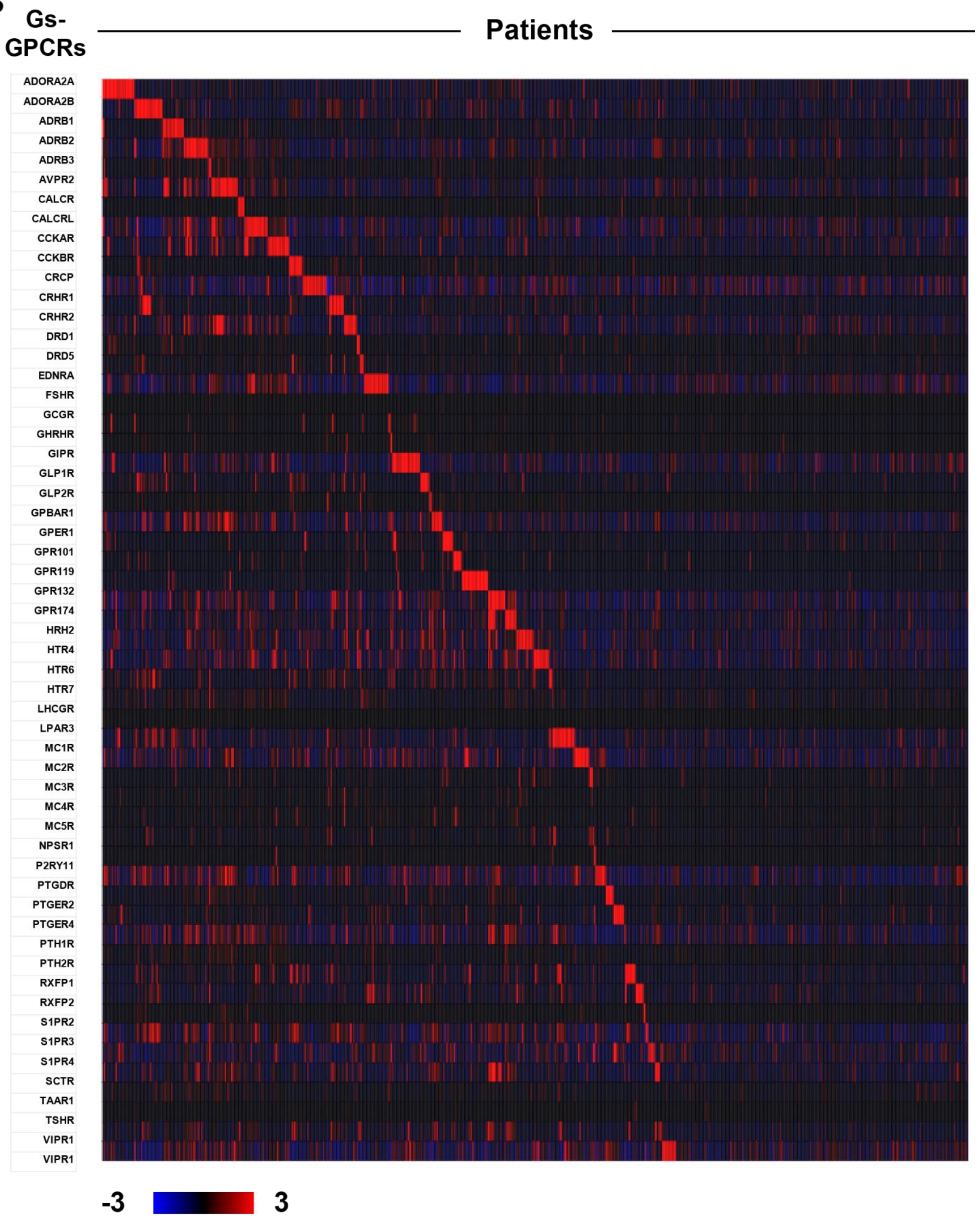
Gs-GPCRs
Patients
ADORA2A
ADORA2B
ADRB1
ADRB2
ADRB3
AVPR2
CALCR
CALCRL
CCKAR
CCKBR
CRCP
CRHR1
CRHR2
DRD1
DRD5
EDNRA
FSHR
GCGR
GHRHR
GIPR
GLP1R
GLP2R
GPBAR1
GPER1
GPR101
GPR119
GPR132
GPR174
HRH2
HTR4
HTR6
HTR7
LHCGR
LPAR3
MC1R
MC2R
MC3R
MC4R
MC5R
NPSR1
P2RY11
PTGDR
PTGER2
PTGER4
PTH1R
PTH2R
RXFP1
RXFP2
S1PR2
S1PR3
S1PR4
SCTR
TAAR1
TSHR
VIPR1
VIPR1
-3
3

C

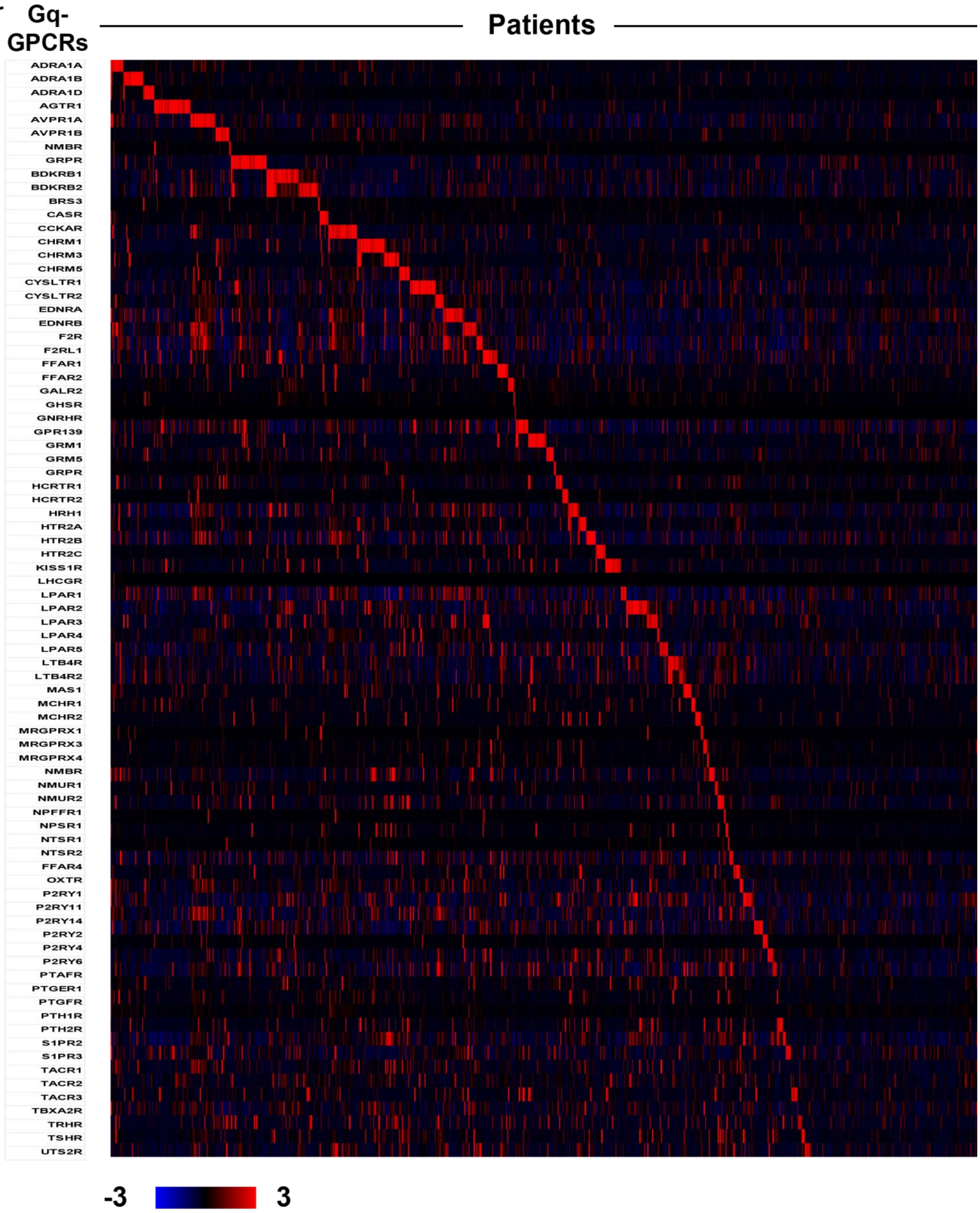
Gq-GPCRs
Patients
ADRA1A
ADRA1B
ADRA1D
AGTR1
AVPR1A
AVPR1B
NMBR
GRPR
BDKRB1
BDKRB2
BRS3
CASR
CCKAR
CHRM1
CHRM3
CHRM5
CYSLTR1
CYSLTR2
EDNRA
EDNRB
F2R
F2RL1
FFAR1
FFAR2
GALR2
GHSR
GNRHR
GPR139
GRM1
GRM5
GRPR
HCRTR1
HCRTR2
HRH1
HTR2A
HTR2B
HTR2C
KISS1R
LHCGR
LPAR1
LPAR2
LPAR3
LPAR4
LPAR5
LTB4R
LTB4R2
MAS1
MCHR1
MCHR2
MRGPRX1
MRGPRX3
MRGPRX4
NMBR
NMUR1
NMUR2
NPFFR1
NPSR1
NTSR1
NTSR2
FFAR4
OXTR
P2RY1
P2RY11
P2RY14
P2RY2
P2RY4
P2RY6
PTAFR
PTGER1
PTGFR
PTH1R
PTH2R
S1PR2
S1PR3
TACR1
TACR2
TACR3
TBXA2R
TRHR
TSHR
UTS2R
-3
3

D

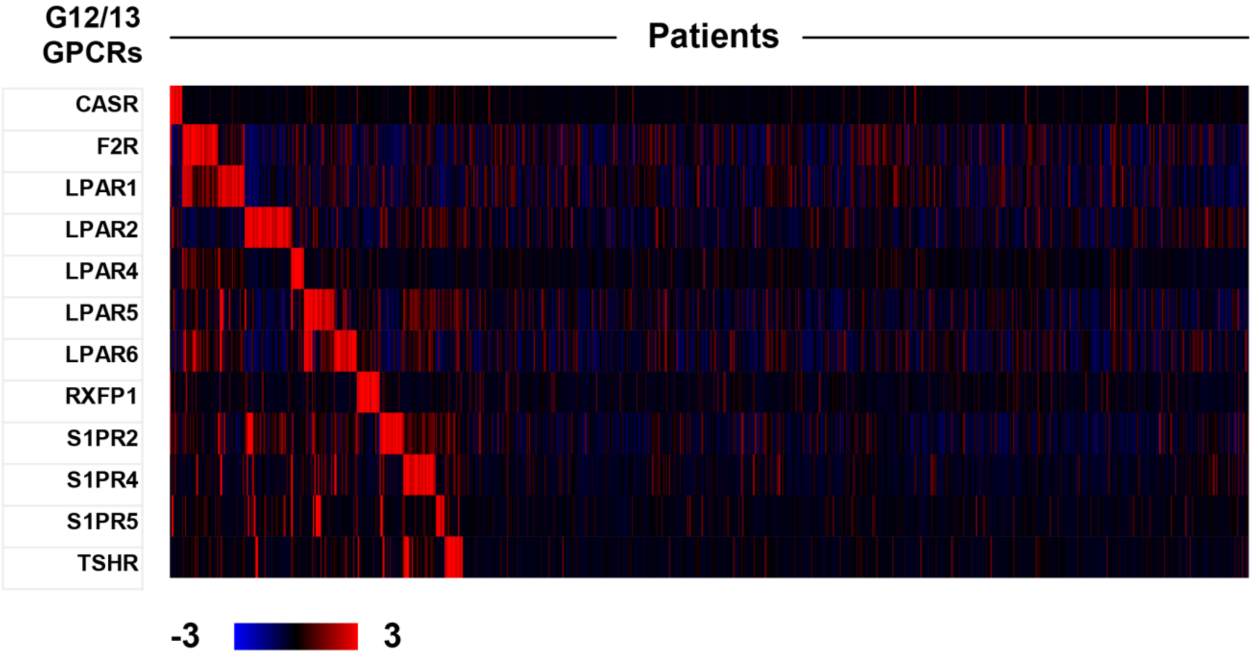
G12/13
GPCRs
Patients
CASR
F2R
LPAR1
LPAR2
LPAR4
LPAR5
LPAR6
RXFP1
S1PR2
S1PR4
S1PR5
TSHR
-3
3

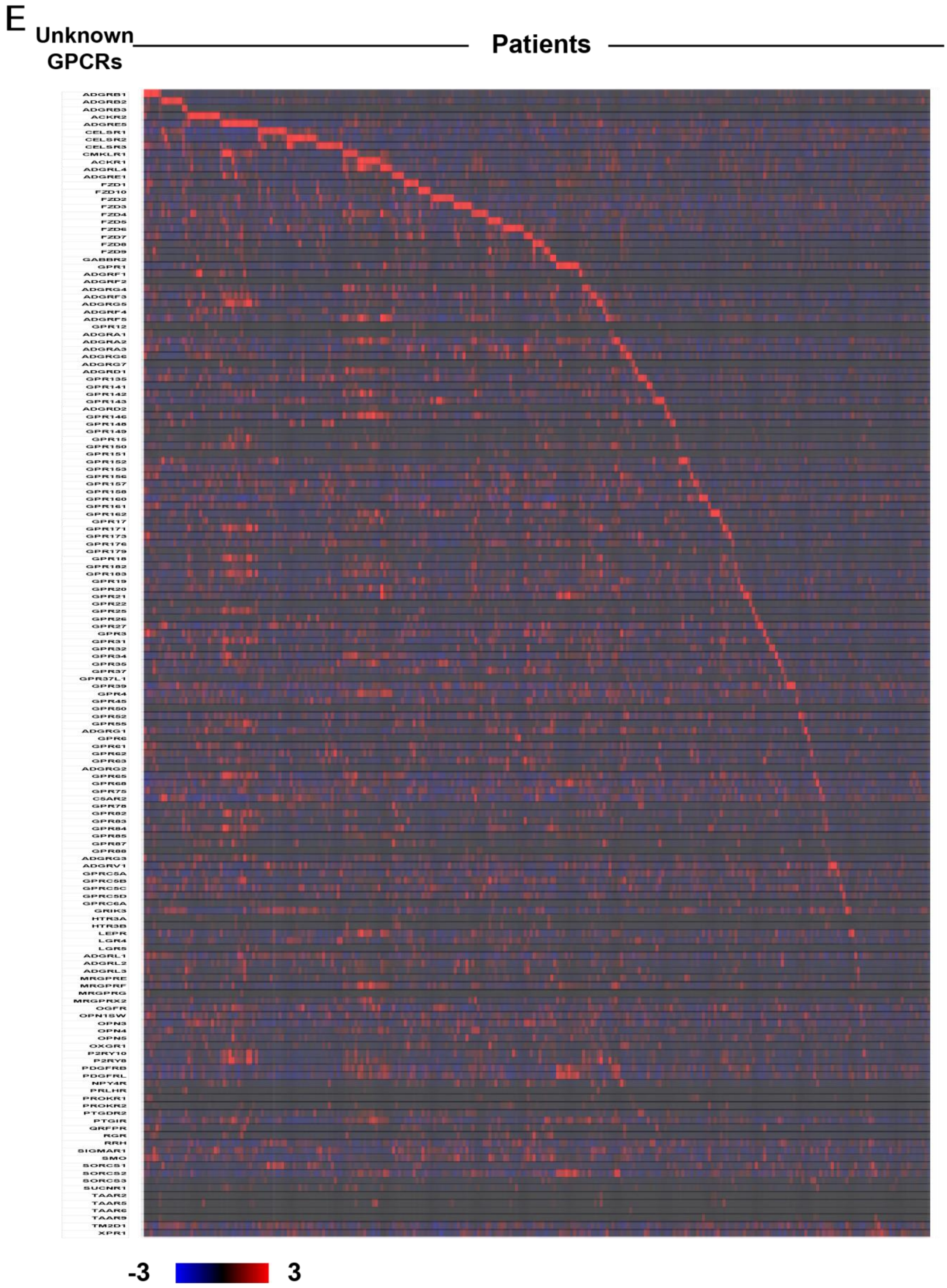


**Supplemental Supplemental Figure 1. Expression of GPCRs in TCGA invasive breast cancer dataset.** A, $G_{i/o}$-coupled GPCRs; B, $G_s$-coupled GPCRs; C, $G_{q/11}$-coupled GPCRs; D, $G_{12/13}$-coupled GPCRs; E, GPCRs with unknown G-protein linkage.

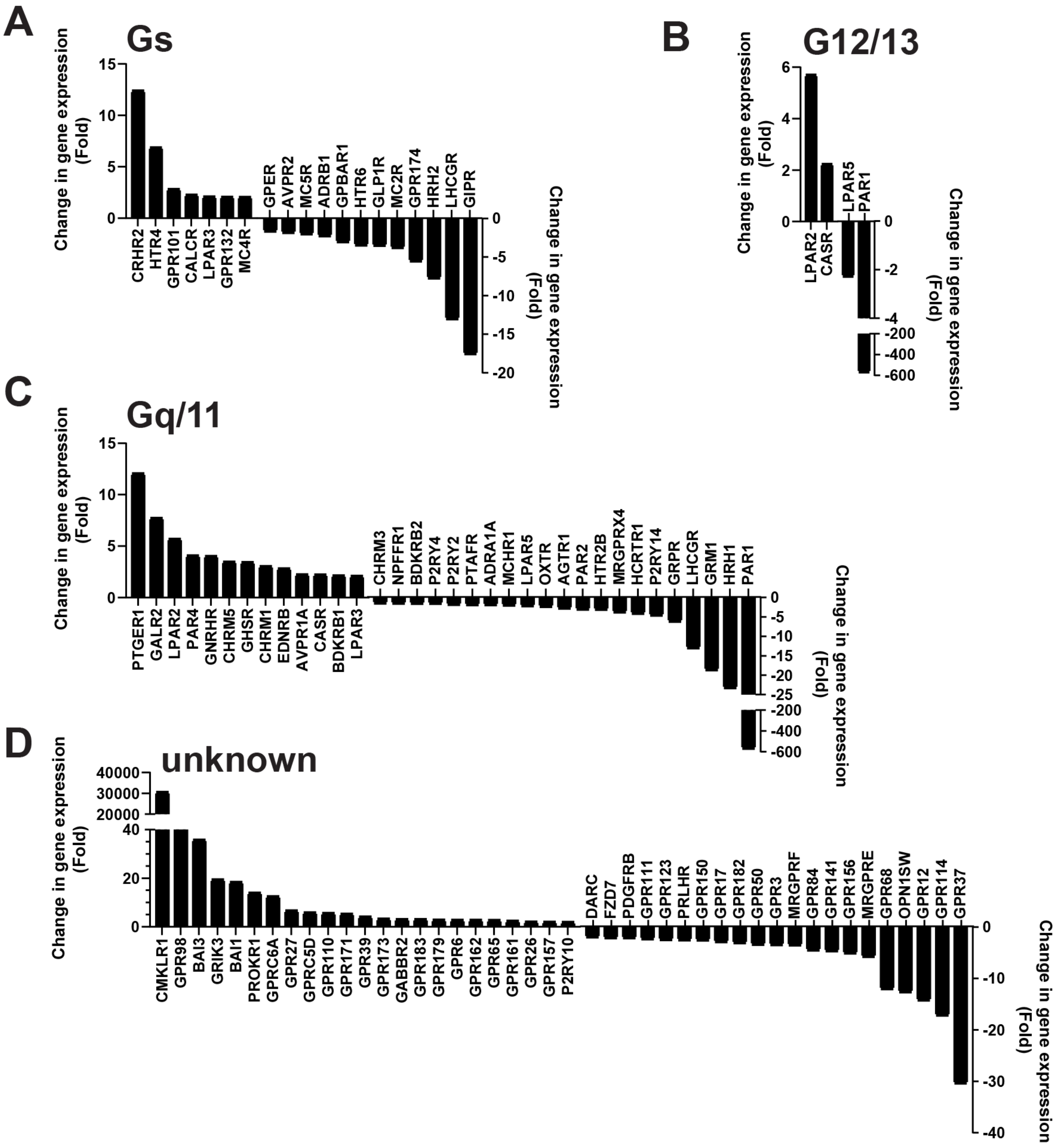


**Figure 2. Altered GPCR expression in Neu cells.** A, $G_s$-coupled GPCRs; B, $G_{q/11}$-coupled GPCRs; C, $G_s$-coupled GPCRs; D, GPCRs with unknown G-protein linkage that show more than 2-fold change of expression in Neu cells as compared to control cells.

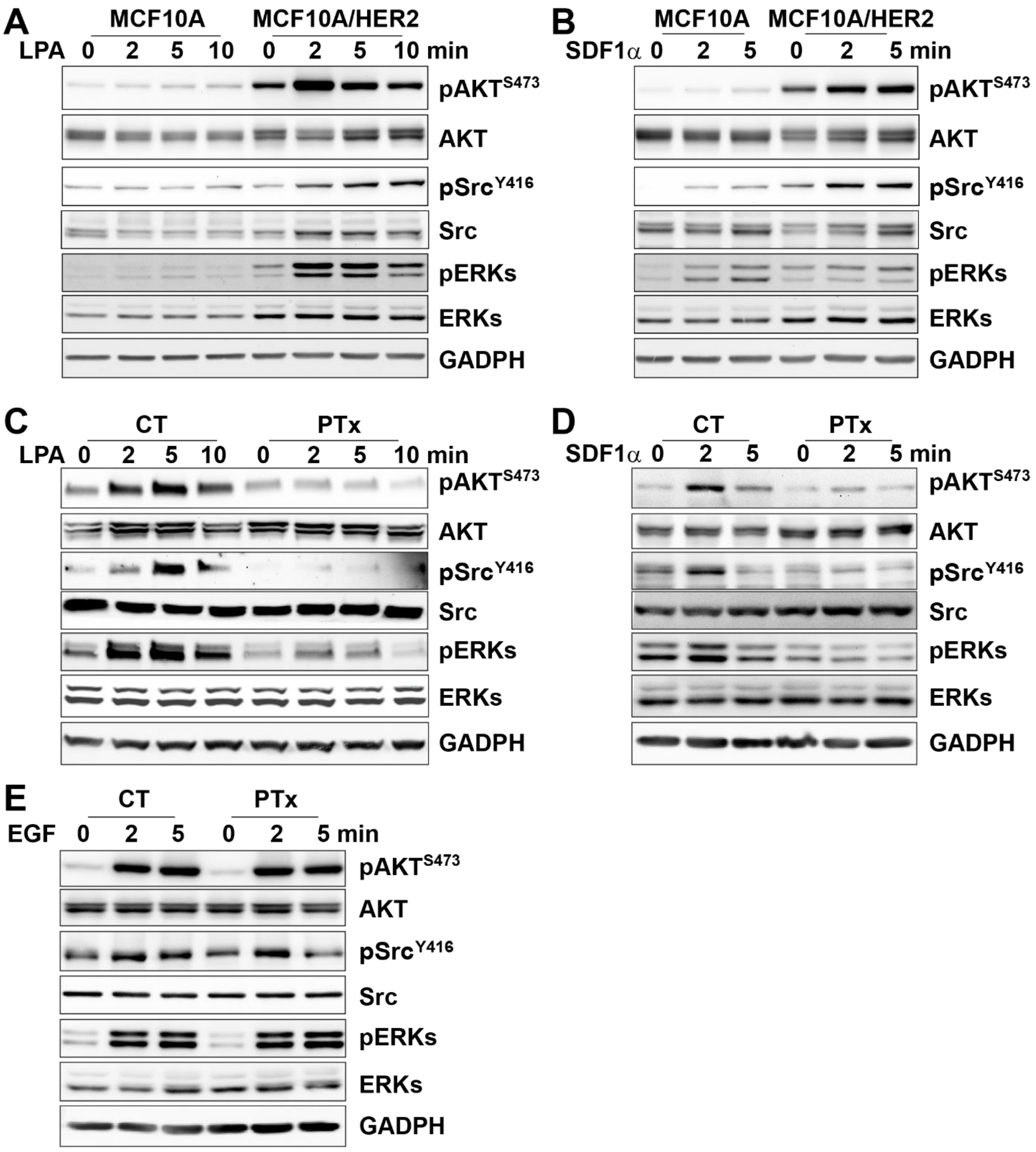


**Supplemental Figure 3. Upregulated Gi/o-GPCR signaling in MCF10A/HER2 cells.** Western blotting showing A-B, the response of MCF10A and MCF10A/HER2 to LPA (A)- and SDF1α (B)-stimulated $AKT^{S473}$ and $Src^{Y416}$ phosphorylation; C-E, the effect of PTx treatment on LPA (C)-, SDF1α (D)- and EGF (E)-stimulated $AKT^{S473}$ and $Src^{Y416}$ phosphorylation in MCF10A/HER2 cells.

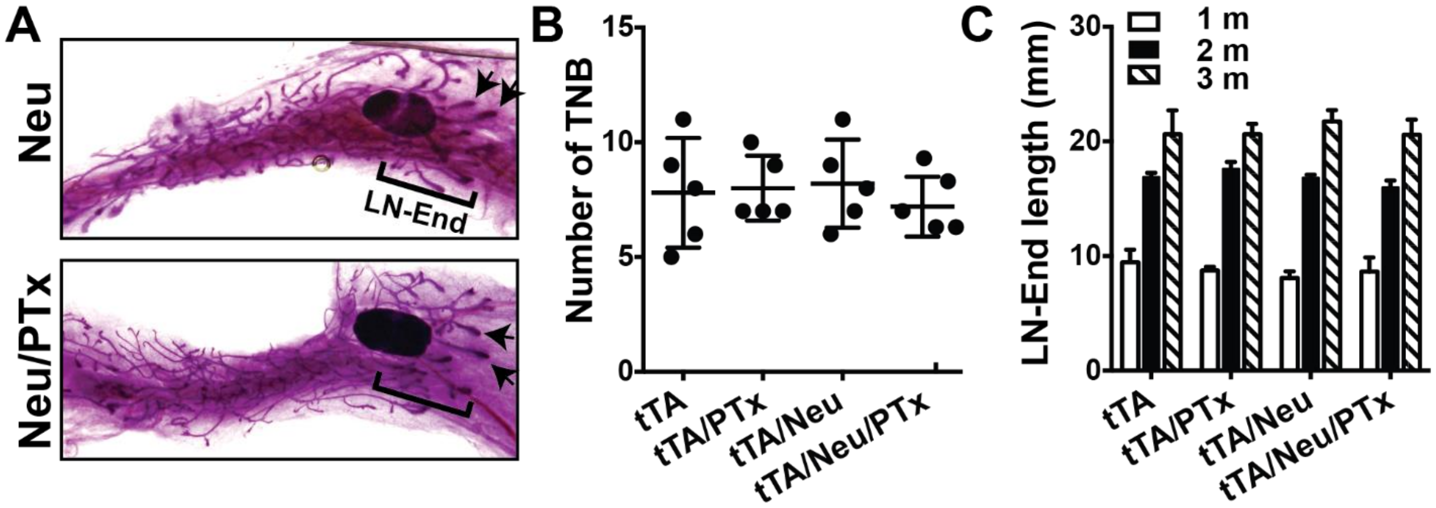


**Supplemental Figure 4. PTx expression does not affect mammary gland development.** A, Whole-mount *in situ* staining showing the terminal end buds (TNBs, indicated by arrows) and length of the ductal distance, measured from the lymph node (LN) to the end of TNBs (LN-End), in the mammary glands from 1-month-old Neu and Neu/PTx mice. B-C, quantitative data showing the number of TNBs (B) and the length of LN-End (C) in mammary glands from transgenic mice at different ages. M: month.

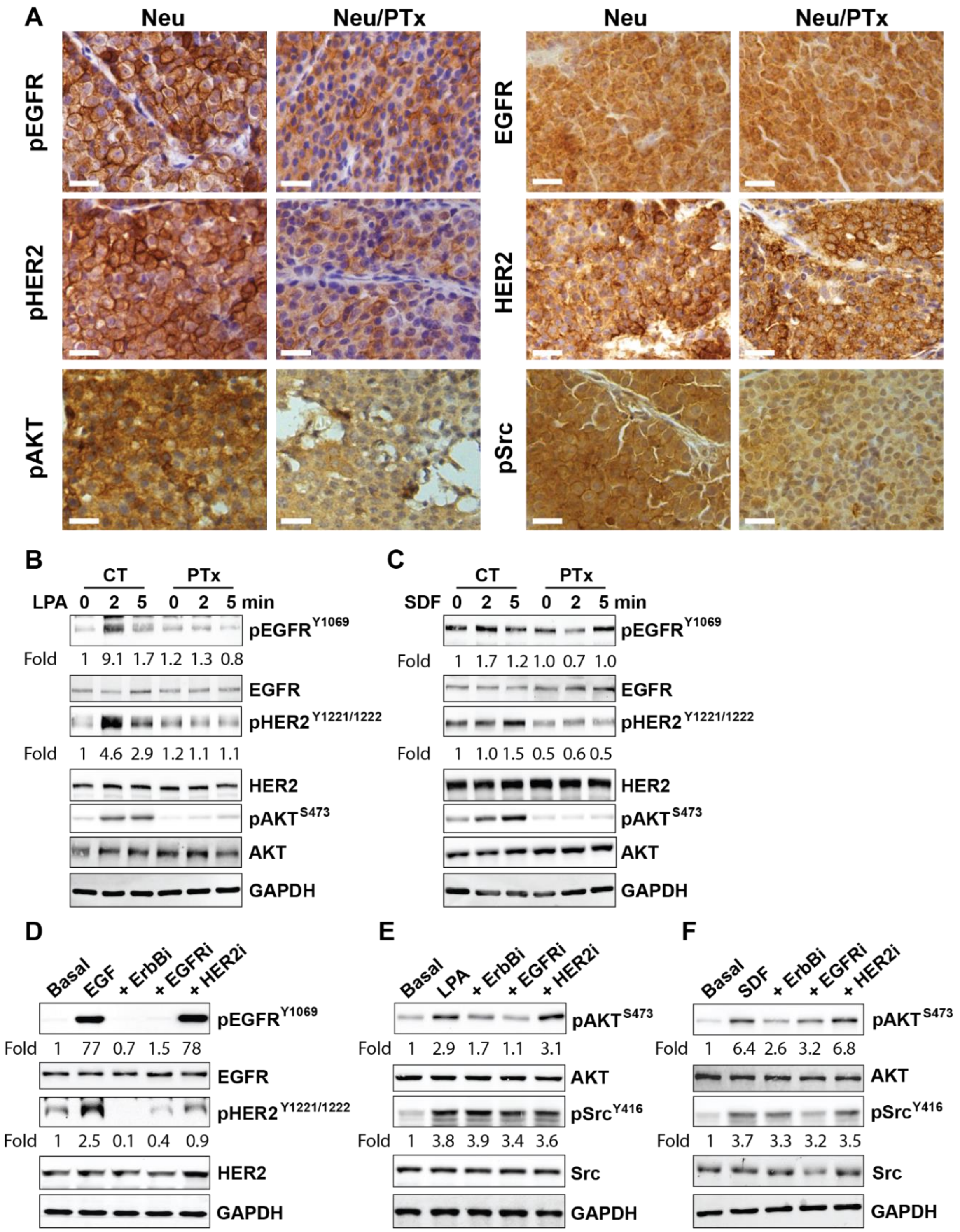


**Supplemental Figure 5. Cross-talk between G$_{i/o}$-GPCRs and EGFR/HER2.** A. IHC staining showing phospho-EGFR$^{Y1069}$ (pEGFR), total EGFR, phospho-HER2$^{Y1221/1222}$ (pHER2), total HER2, phospho-AKT$^{S473}$ (pAKT), and phospho-Src$^{Y416}$ (pSrc), in Neu and Neu/PTx tumors. Scale bar=10 μm. B-C, Western blotting showing phosphorylation of EGFR, HER2 and AKT in MCF10A/HER2 cells stimulated with LPA (B) and SDF1α (C) and treated with vehicle (CT) or PTx. D-F, the effect of the pan-ErbB-, EGFR- and HER2-selective inhibitors on the activation of EGFR and HER2 by EGF in Neu cells (D) and the phosphorylation of AKT$^{S473}$ and Src$^{Y416}$ by LPA (E) and SDF1 α (F) stimulation in MCF10A/HER2 cells. The phosphorylation of EGFR$^{Y1068}$, HER2$^{Y1221/1222}$, AKT$^{S473}$ and Src$^{Y416}$ was quantified as the ratio of the phosphorylated to total proteins and expressed as the fold increase over the basal, which is indicated underneath the images.
